# Citrus photosynthesis and morphology acclimate to phloem-affecting huanglongbing disease at the leaf and shoot levels

**DOI:** 10.1101/2021.07.13.452140

**Authors:** Mark Keeley, Diane Rowland, Christopher Vincent

## Abstract

Huanglongbing (HLB) is a phloem-affecting disease of citrus that reduces growth and has impacted global citrus production. HLB is caused by a phloem-limited bacterium (*Candidatus* Liberibacter asiaticus; *C*Las). By inhibiting phloem function, HLB stunts sink growth, including reducing production of new shoots and leaves, and induces hyperaccumulation of foliar starch. HLB induces feedback inhibition of photosynthesis by reducing foliar carbohydrate export. In this work we assessed the relationship of bacterial distribution within the foliage, foliar starch accumulation, and net CO_2_ assimilation (*A*_net_). Because HLB impacts canopy morphology, we developed a chamber to measure whole-shoot *A*_net_ to test the effects of HLB at both leaf and shoot levels. Whole-shoot-level *A*_net_ saturated at high irradiance, and green stems had high photosynthetic rates compared to leaves. Starch accumulation was correlated with bacterial population, and starch was negatively correlated with *A*_net_ at the leaf level but not at the shoot level. Starch increased initially after infection, then decreased progressively with increasing length of infection. HLB infection reduced *A*_net_ at the leaf level, but increased it at the whole shoot level, in association with reduced leaf size and greater relative contribution of stems to photosynthetic surface area. Although HLB-increased photosynthetic efficiency, total carbon fixed per shoot decreased because photosynthetic surface area was reduced. We conclude that the localized effects of infection on photosynthesis are mitigated by whole shoot morphological acclimation over time. Stems contribute important proportions of whole shoot *A*_net_, and these contributions are likely increased by the morphological acclimation induced by HLB.

## Introduction

Huanglongbing (HLB) is a disease of citrus causing economic loss to the citrus industry around the world (Bové, 2006; Singerman and Useche, 2016). Its causal agent, *Candidatus* Liberibacter asiaticus (*C*Las), vectored by the Asian citrus psyllid (ACP; *Diaphorina citri* Kuwayama), is a phloem-limited bacterium which effects phloem dysfunction leading to starch accumulation in leaves and loss of sink growth (Achor et al., 2020; Bové, 2006), as well as eventual development of thin canopies (Tang et al., 2019).

Because of the reduction in phloem function, HLB is hypothesized to lead to feedback inhibition of photosynthesis (Cimò et al. 2013). Assessing chlorophyll fluorescence, a reduction in maximal PSII quantum yield was found in *C*Las-infected citrus leaves compared to uninfected leaves (Cen et al., 2017; Sagaram & Burns, 2009). On the one hand, starch accumulation in the leaves has been noted as a symptom of HLB, which indicates a surplus of fixation relative to export (Etxeberria et al., 2009; Folimonova & Achor, 2010; Schneider, 1968). Starch accumulation can disintegrate the chloroplast membrane (Nebauer et al., 2011; Snyder-Leiby & Wang, 2008), which may contribute to the reduction in PSII yield noted above. However, no measures of the impacts of HLB on net CO_2_ assimilation (*A*_net_) have been reported.

Measures of *A*_net_ in citrus have previously been taken at the leaf level (Kriedemann, 1968; Mesquita et al., 2016; Tsagarakis et al., 2012; Vu and Yelenosky, 1988; Vu et al., 1986; Zambrosi et al., 2011) and at the field level (Contreras et al. 2017; Er-Raki et al., 2009; Maestre-Valero et al., 2017; Penddinti and Kambhammettu, 2019). Due to the specificity of the leaf level measurements (light levels, angle of radiation incidence, leaf position within the canopy), leaf-level data is difficult to extrapolate to the whole canopy. Because HLB induces reductions in canopy density (Tang et al. 2019), larger scale measures may be more indicative of changes in carbon assimilation than leaf-level measures.

In this study, our objective was to compare relationships of *A*_net_, *C*Las infection, and foliar symptoms at the leaf and shoot level. To achieve our objective, we first developed and validated the use of a gas exchange chamber capable of measuring whole shoots. We assessed the value of such measures by comparing leaf-level and shoot-level light response curves. In our subsequent HLB experiment, we hypothesized that: 1) greater time of *C*Las inoculation exposure would increase starch content; 2) higher starch content, as an indicator of reduced leaf carbohydrate export, would correlate with downregulation of *A*_net_; and 3) because HLB causes morphological changes at the canopy level, mean leaf area, leaf number, and stem surface area and length were expected to affect *A*_net_ at the shoot level.

## Materials and Methods

### Study 1: Whole shoot gas exchange chamber development and testing

#### Plant Material

The trees utilized were greenhouse-raised, four-year-old ‘Valencia’ sweet orange (*Citrus x sinensis*) on ‘Kuharske’ (*C. sinensis* x *Poncirus trifoliata* (L.) Raf.) rootstock in 20-L pots. Plants were maintained with recommended rates of slow-release fertilizer and 3x weekly irrigation with sufficient volume to exceed field capacity of the pots.

#### Chamber construction

The chamber was built of acrylic tubing, acrylic glue, and a 3-D printed adapter designed and provided by Jason Hupp, LI-COR Bioscience, Inc. Lincoln, NE, USA. The chamber was fit to an infrared gas analyzer (LI-6800, LI-COR Bioscience, Inc. Lincoln, NE, USA) using a custom chamber adapter (LI-6800-19, LI-COR Bioscience, Inc. Lincoln, NE, USA). To seal the opening, a large conical stopper was step-drilled and cut to provide an easy-to-fit cap around the stem of the branch (Appendix S1). The chamber could then be made airtight using plumbers’ putty.

Leaves and shoots were measured on trees recently acclimated to the relative darkness of the laboratory environment (2-4 μmol photons m^-2^ s^-1^). Leaf-level measurements were recorded for 15 minutes on three leaves at photosynthetic photon flux density (PPFD) of 1000 μmols m^-2^ s^-1^. Environmental settings for the leaf level were set at flow 300 μmol s^-1^, 0.2 kPa pressure, 60% relative humidity, reference CO_2_ at 400 µmol CO_2_ mol^-1^, and a fan speed of 10,000 RPM. Chamber measurements were logged every 15 seconds for five minutes after a 15-minute stabilization period under controlled conditions at PPFD of 1000 μmol m^-2^ s^-1^. Environmental settings for the whole-shoot chamber were set based on volume, 1000 cubic centimeters, at flow 1000 μmol s^-1^, 0.2 kPa pressure, 60% relative humidity, reference CO_2_ of 400 µmol CO_2_ mol^-1^, and a fan speed of 14,000 RPM to maximize air mixture. Increases in air flow rate from 300 μmol s^-1^ in leaf measurements to 1000 μmol s^-1^ in whole -shoot chamber compensated for the increase in photosynthetic material and increased volume present in the custom chamber. The fan speed was increased to 14,000 RPM to help mix air in the chamber as recommended by the IRGA manufacturer. Using the red-light capability of the red, far red, and blue light adjustable fluorometer head unit of the LI-6800F, light intensity was controlled for the leaf-level measurements. An LED Grow Light of similar spectral composition (ZXMean 1500W Full Spectrum LED, BliGli, Wuhan HuBei, China) was manipulated by distance and a dimming switch (Credenza Plug-In Lamp CFL-LED Dimmer, Lutron Electronics Co., Coopersburg, PA, USA) to achieve desired PPFD for the whole-shoot chamber. PPFD for whole-shoot measurements was measured using the quantum sensor of the LI-6800 at the mid-point of the shoot adjacent to the whole-shoot chamber.

#### Stem assimilation

To determine stem contribution to *A*_net_ of the shoot (the leaf and stem material together), the stems were covered with fitted black drinking straws following the method of Nilsen and Bao (1990) testing stem contributions to photosynthesis in soybean (*Glycine max* L.). Shoots were exposed to PPFD 1400 μmol m^-2^ s^-1^ as described above. The shoot was then removed from the chamber and black straws were measured to fit the space between petioles on the stem, sliced lengthwise and wrapped around the stem (Appendix S2). Gas exchange measurements then rerecorded with the darkened stems. The LED Grow Light was then turned off and the custom chamber was covered by a black cloth to achieve 0 μmol m^-2^ s^-1^ PPFD to determine respiration of the shoot, again following the same stabilization and logging duration. Measurements used were: whole shoot in the light (Shoot *A*_net_), lit shoot with only the stem darkened stem, and shoot dark respiration *R*_d_, measured as net CO_2_ exchange at 0 μmol m^-2^ s^-1^ PPFD (shoot respiration) were used for analysis. Past measurements of whole-plant photosynthesis have considered gross photosynthesis to be *A*_net_ – *R*_d_ (Hwang and Morris 1994), other authors have pointed out that gross photosynthesis is more accurately interpreted as *A*_net_ – *R*_d_ – photorespiration, and that *A*_net_ – *R*_d_ is better described as apparent photosynthesis (*A*_app_) (Wohlfahrt and Gu 2015). Measurements of *A* under the different measurement conditions were used to calculate *A*_app_ contributions of the stem and leaves as follows:

- Shoot *A*_net_: shoot net photosynthesis, measured as *A* of shoot at 1400 µmol m^-2^ s^-1^ PPFD; inherently the sum of stem and leaf *A*_app_ minus respiration (shoot *R*_d_).
- Stem *A*_app_: stem apparent *A*; measured as the difference between shoot *A*_app_ under full illumination and *A*_app_ with darkened stem.
- Shoot *R*: whole shoot respiration, measured as |*A*| at 0 µmol m^-2^ s^-1^ PPFD.
- Shoot *A*_app_: shoot apparent assimilation, estimated as shoot *A*_net_ – shoot *R*_d_.
- Leaf *A*_app_: leaf apparent assimilation of all leaves on the shoot, estimated as shoot *A*_app_ – stem *A*_app_

Immediately following gas exchange measurements, shoots were excised and defoliated for leaf area analysis using the LI-3100 (LI-COR Biosciences, Inc. Lincoln, NE, USA). Stems were measured for length and diameter (Appendix S3) to calculate surface area using an adjusted formula for the lateral area of a truncated cone (Equation 1). Due to the irregular shape of citrus stems, two diameter measurements were taken at the widest and narrowest widths at the base, middle, and tip of the stem. The circumference of an ellipse is approximately equal to that of a circle with a radius of the average of the widest and narrowest point of the ellipse.

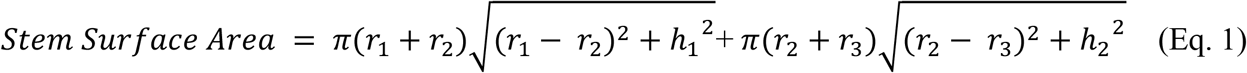

Where:

r_1_= radius of stem base

r_2_= radius of stem middle

r_3_= radius of stem tip

h_1_ =length between measurement point of r_1_ and r_2_

h_2_ =length between measurement point of r_2_ and r_3_

Leaves and stems were dried at 60 °C for 72 hours and weighed separately. Leaf and stem surface area and dry weight were used to calculate *A*_app_ rates per meter squared and per gram dry weight.

#### Leaf and shoot light response curves

Nine fully mature young shoots, three from each of the same trees were measured in the laboratory, because ambient light in the greenhouse prevented low-light manipulation of shoot level environment. Shoots were placed in the shoot gas exchange chamber and carbon assimilation was measured at PPFD of 0, 25, 50, 100, 150, 200, 400, 600, 800, 1000, 1400, and 1800 μmols m^-2^ s^-1^. Environmental settings, stability criteria, and logging were the same as previous whole-shoot measurements (Appendix S4). The largest, healthiest leaf (assessed visually) on each of the nine measured shoots was selected to measure a leaf-level comparison (Appendix S5) measurement at PPFD of 0, 25, 50, 100, 150, 200, 400, 600, 800, 1000, 1400, and 1800 μmols m^-2^ s^-1^. Environmental settings and stability criteria were set as in the previously described measurements.

### Study 2: HLB impacts on leaf and shoot net CO_2_ assimilation

#### Plant material selection

Trees were selected based on length of exposure to citrus greening infection. The trees utilized were greenhouse-raised, four to five-year-old sweet orange ‘Valencia’ on ‘Kuharske’ rootstock in 20-L pots. These trees had been exposed to ACP inoculation of *C*Las by growing plants in infested screenhouses for varying lengths of time. Exposure categories were defined as follows: the High category of exposure had been exposed to infected ACP feeding and breeding for 2.5 years; the Mid category of exposure had been exposed to infected ACP feeding and breeding for 1.5 years; and the Low category of exposure had been exposed to infected ACP feeding and breeding for 1 year. The uninfected trees had not been exposed to infected ACP. Three trees of each category were arranged in a greenhouse in a randomized complete block design, totaling 12 trees for the study.

Shoots and leaves were selected on each tree for measurements. Three shoots per tree were selected from the exterior of the canopy. Shoots were chosen based on accessibility of the instrument using the custom chamber, and only those with at least three leaves suitable for leaf measurement using the standard leaf chamber were selected. Thirty-six shoots were measured in this study (3 shoots x 3 trees x 4 exposure groups).Three leaves were selected per shoot. These were distributed along the length of the shoot, always using one apical leaf. To be selected, leaves had to be large enough to completely cover the leaf chamber (6-cm^2^ circle). A total of 108 leaves were measured for this study (36 shoots x 3 leaves).

#### Gas exchange measurements

*A*_net_, transpiration (*E*), and conductance were measured in using the LI-6800 (LI-COR Bioscience, Inc. Lincoln, NE, USA). In the case of leaf-level measurements, conductance represented stomatal conductance to water (*g*_*s*_) while the shoot level measurements estimate shoot conductance to water (shoot *g*), because non-stomatal pores, such as lenticels may contribute significantly conductance at the shoot level. Leaf chamber environmental settings were set at a PPFD of 1000 µmol m^-2^ s^-1^ and shoot chamber was placed at PPFD of 1400 µmol m^-2^ s^-1^ with all other settings according to those described above for Study 1 for leaf and shoot gas exchange measurements.

#### CLas quantification

Midrib and petiole tissues excised from selected leaves were chopped using a razor blade and bead-beaten using Vortex Genie 2 (Scientific Industries Bohemia, NY, USA) to a fine homogenized powder in liquid nitrogen. Total genomic DNA was extracted and purified from 100 mg of homogenized material using GeneJET Plant Genomic DNA Purification Mini Kits (Thermo Fisher Scientific Waltham, MA, USA). Samples were processed following manufacturer instructions and stored at −20° C. Primers and TaqMan probes were obtained for *C*Las based on method developed by Coy et al. (2014). The PCR master mix was used with 10 μL TaqMan Buffer, 8.6 μL sterile deionized water, 0.4 μL CQUL primer, and 1 μL of extracted purified DNA. PCR reactions performed with standards using QuantStudio 3 Real-time PCR (Applied Biosystems-Thermo Fisher Scientific Waltham, MA, USA) yielded cycle threshold values (Ct). Bacterial copy-count estimates were generated by regression using processed Ct results from standards of known copy quantity, processed in each plate batch. At the shoot level Ct and bacterial copy number were considered to be the mean of the measurements in the three selected leaves on each shoot.

#### Starch content

Starch content was determined by extracting starch from a single 28.27 mm^2^ hole punch on the selected leaf blade (Whitaker et al., 2014). The leaf punch was homogenized for 2.5 minutes at 6500 RPM in 500 μL of distilled water with five beads. Samples were boiled for 10 minutes at 200° C and agitated prior to a two-minute centrifuge at 2500 rpm, after which 300 μL of the supernatant was added to 900 μL pure ethanol and agitated for three seconds to precipitate starch. Prepared samples were centrifuged at 14000 RPM for 10 minutes and the supernatant was discarded and replaced by one mL of deionized water. Samples were then shaken for four-minutes to dissolve the starch pellet and two 250 μL of prepared sample were each placed in individual wells of a 96 well microplate, and 50 μL of 2% iodine was added to each well. Absorbance at 595nm was read using SpectraMax 250 microplate reader (Molecular Devices San Jose, CA, USA). Linear functions based on standard curve concentrations included in each plate were generated with the absorbance of standards to estimate starch content. As for *C*Las variables, at the shoot level starch values were considered to be the mean of the measurements of the three selected leaves on each shoot.

#### Visual rating

A rating of the visual symptoms of *C*Las infection was assessed on a scale of 0-10, where 0 = no symptom, 10= full symptom expression, on each leaf. Prior to assessment a scale of leaf symptoms was generated using photos of sampled *C*Las infected leaves to ensure consistency in ratings. All disease ratings were performed by the same individual. Visual symptom ratings used in the leaf level measurements were recorded for every leaf on the measured shoot; an average of the assessment of all leaves was used for the shoot value.

#### Shoot morphology

Total leaf area per shoot was measured using a LI-3100 (LI-COR Bioscience, Inc. Lincoln, NE, USA) immediately following the assimilation measurements and visual ratings, prior to sampling for qPCR and starch content. Stem length and diameter in two directions was measured using a micrometer to generate a stem surface area estimates using Eq. 3. Average internode length was calculated by dividing the stem length by the total number of leaves per stem.

#### Analysis

Analysis of variance was performed using the anova() command in base R, after assessing appropriateness of parametric tests by testing for normal distribution, to assess the effect of treatment groups (R Core Team, 2019, Version 3.6.1). Where exposure group was significant, Fisher’s LSD was used as a post hoc test for least significant difference between the four tree categories used for all variables using the {agricolae} package (de Mendiburu, 2020). Correlation analysis was generated among the variables at the shoot and leaf-level and correlation matrices were tested for significance of Pearson’s correlations using the {Hmisc}package (Harrell, 2020). For correlation analyses, because some variables might co-vary in the case of disease, tests were performed with and without the HLB-plants. For variables with significant correlations, linear regression models were generated at both shoot and leaf-level using the lm() command, similarly comparing relationships with and without HLB-plants. The models were tested for significant relationships using anova(). All linear models nested shoot within tree, and leaf-level data was nested leaf within shoot. A natural log transformation of bacterial copy count and starch content facilitated a linear relationship between the transformed data. Light response curves for leaf and shoot were fit using an nonlinear least squares regression using the self-starting asymptotic function in R (Pinheiro et al., 2014) with a function supplied by Tomeo (2017). This function fits an asymptotic function and calculates the parameters of fully saturated photosynthesis (*A*_sat_), quantum yield of photosynthesis (Φ), dark respiration (*R*_d_), light compensation point (LCP), and PPFD of 75% saturation (Q_sat75_). These parameters were then subjected to analysis of variance to test the effects of shoot vs. leaf light responses.

## Results

### Stem contributes significantly to whole-shoot photosynthesis

Vegetative stem on young shoots contributed 5.7% to total shoot surface area, 17.0% to total shoot mass, and 10.7% to total shoot photosynthesis (μmol CO_2_ s^-1^; Table 1). Mean stem *A*_app_ was nearly double leaf *A*_app_ (9.62 μmol CO_2_ m^-2^ s^-1^ and 5.13 μmol CO_2_ m^-2^ s^-1^, respectively; Figure 2).

**Table 1.** Percent contribution of leaves and stems to the shoot surface area and apparent CO_2_ assimilation (*A*_app_) in ‘Valencia’ sweet orange (*Citrus x sinensis* [L.] Osbeck).

| Measurement basis | Leaf | Stem |
| --- | --- | --- |
| Surface area | 94.3 ± 1.6 % | 5.6 ± 1.6 % |
| Mass | 83.0 ± 4.8% | 17.0 ± 4.8% |
| Per shoot $A_{app}$ | 89.2 ± 8.5 % | 10.7 ± 8.5% |

**Figure 1.**
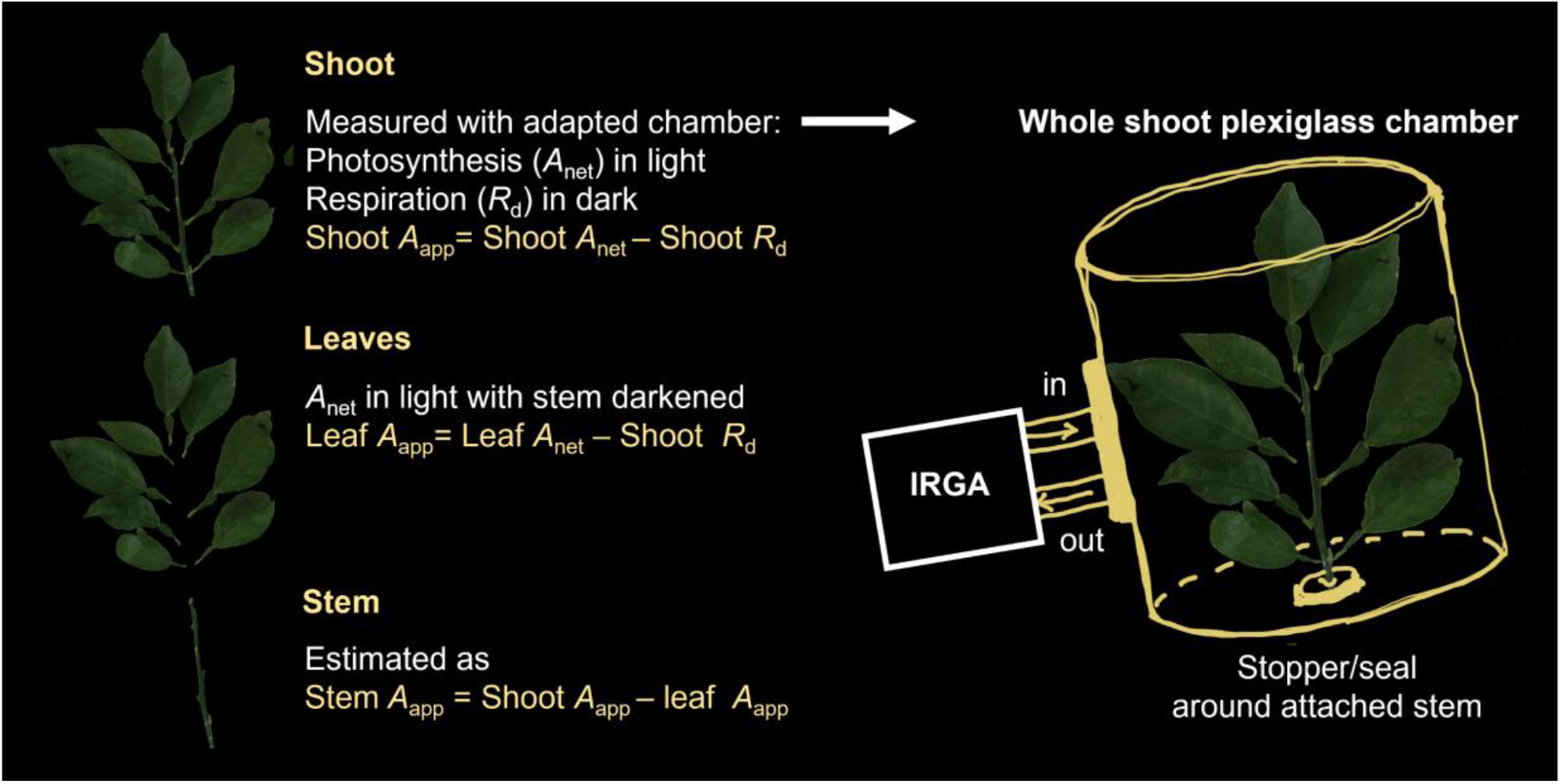
Illustration of the use of a whole-shoot chamber to estimate apparent photosynthesis (*A*_app_) of whole shoots, leaves, and stem.

**Figure 2.**
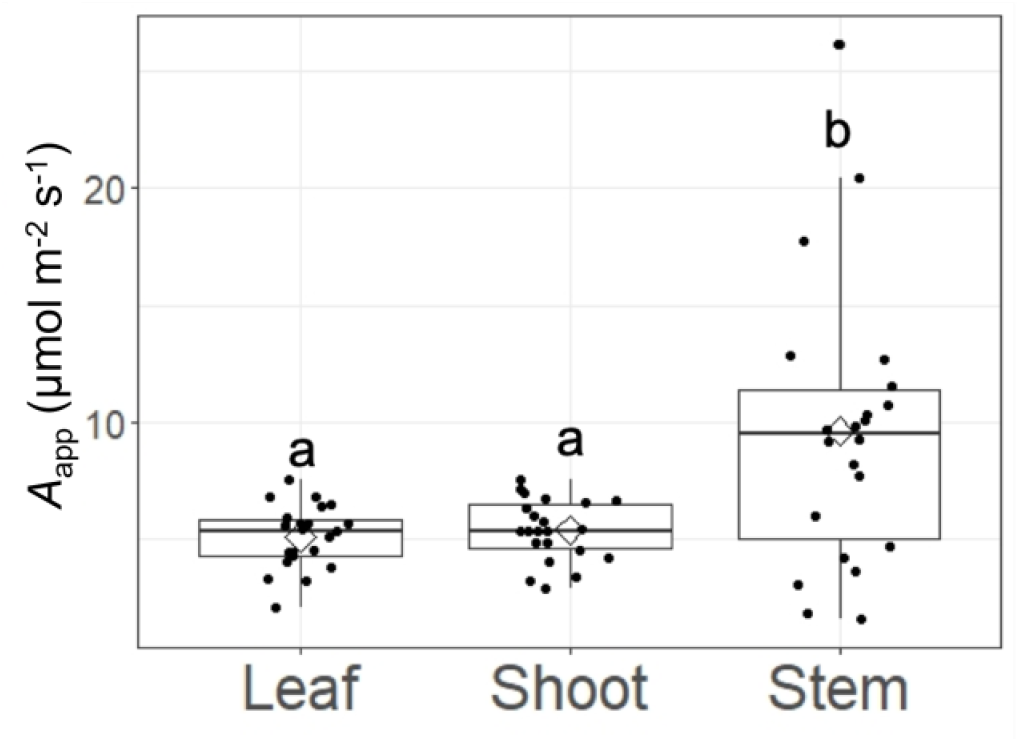
Apparent assimilation (net assimilation – dark respiration) for ‘Valencia’ sweet orange (*Citrus x sinensis* [L.] Osbeck) leaf, stem and shoot. Letters within each plot denote means that are significantly different at P≤0.05 according to Fisher’s least significant differences.

### Light response curves differ between shoots and leaves

Leaves and shoots had distinct light response curves with differences in all parameters tested, except for quantum yield (Table 2). Leaf level *R*_d_ was more than 3x that of the shoot level *R*_d_, resulting in a correspondingly higher LCP, despite the less acute rise in *A*_net_ response to PPFD (Figure 3). Leaf level 75% saturation was achieved at approximately 528 μmol photons m^-2^ s^-1^, while that of shoots was estimated at 1532 μmol m^-2^ s^-1^. The corresponding *A*_sat_ was also approximately 20% higher in shoots than in leaves.

**Table 2.** Analysis of variance and means comparison of parameters of a light response curve function fit to leaf and whole shoot light response curves of ‘Valencia’ sweet orange (*Citrus x sinensis* [L.] Osbeck). Values are means ± standard error. Values followed by different letters are significantly different at P<0.05. *A*_sat_: light-saturated net CO_2_ assimilation (µmol CO_2_ m^-2^ s^-1^); Φ:quantum yield (mols CO_2_ fixed:mol photons absorbed); *R*_d_: dark respiration (µmol CO_2_ m^-2^ s^- 1^); LCP: light compensation point (µmol photosynthetically active radiation m^-2^ s^-1^); Qsat75: photosynthetic flux density at 75% saturation (µmol photosynthetically active radiation m^-2^ s^-1^).

| Function parameter | Leaf | Shoot | F-value | P-value |
| --- | --- | --- | --- | --- |
| $A_{sat}$ | 7.7 ± 0.46 b | 10.4 ± 0.74 a | 8.7 | 0.01 |
| $\Phi$ | 0.044 ± 0.002 | 0.053 ± 0.02 | 0.2 | 0.65 |
| $R_d$ | 1.2 ± 0.12 a | 0.35 ± 0.04 b | 42.4 | <0.0001 |
| LCP | 28.0 ± 8.4 a | 14.7 ± 10.8 b | 7.9 | 0.01 |
| $Q_{sat75}$ | 528 ± 80 b | 1532 ± 226 a | 17.3 | 0.0016 |

**Figure 3.**
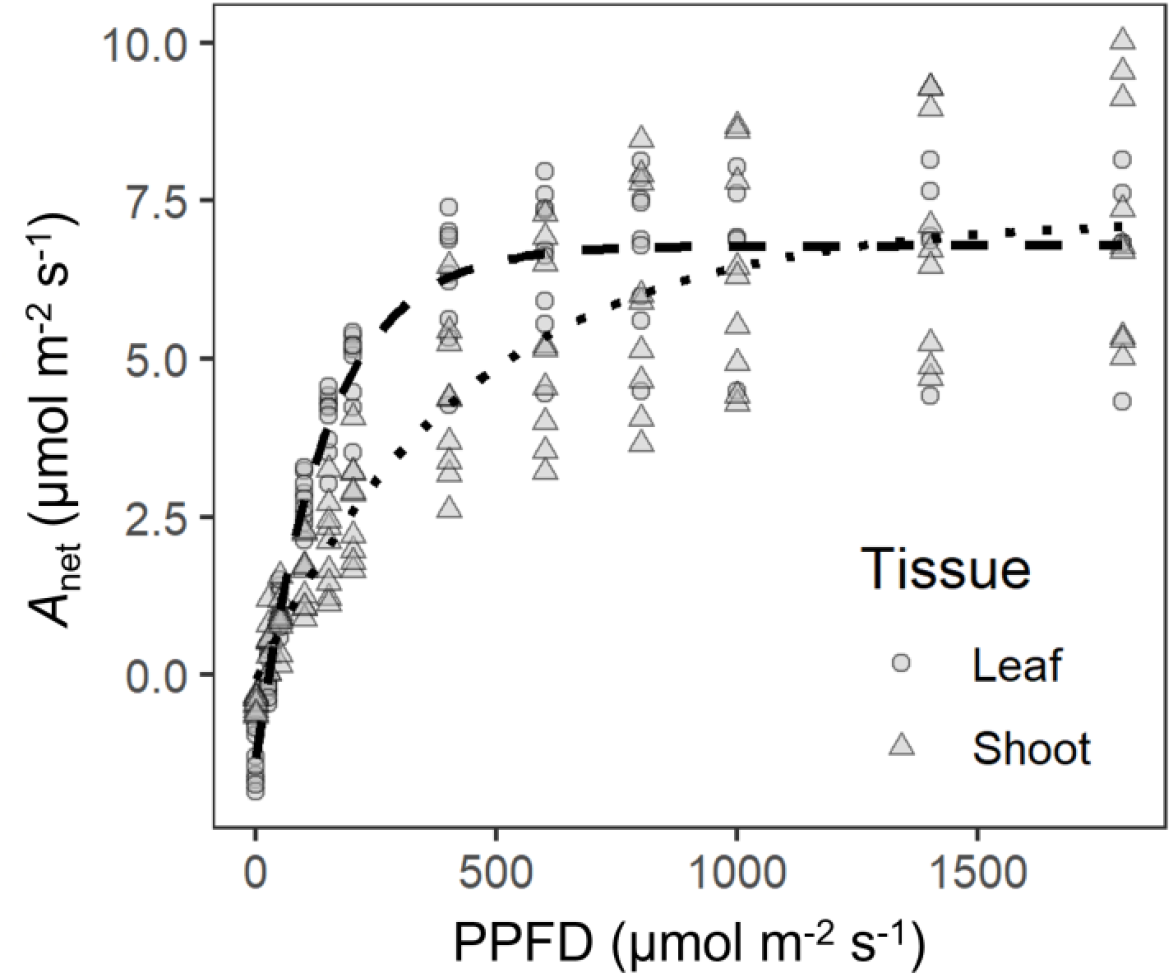
Distinct carbon assimilation light response curves of ‘Valencia’ sweet orange (*Citrus x sinensis* [L.] Osbeck) leaf (dashed line) and shoot (dotted line). All points are biological replicates.

### Photosynthesis and starch are affected by CLas exposure

Uninfected leaves had a higher *A*_net_ (mean of 3.97 μmol CO_2_ m^-2^ s^-1^) than infected leaves; however, as *C*Las exposure increased from low, to medium, to high so did *A*_net_ (means of 1.91, 2.61, 2.70 μmol CO_2_ m^-2^ s^-1^, respectively; Table 3). Leaf-level *g*_*sw*_ intermediate in the uninfected category, while it was lower in the low and medium exposure groups, while *g*_*sw*_ was highest in the high exposure group. A similar pattern was observed for leaf-level *E*, though the *P*-value for the effect of exposure category was inconclusive at *P*=0.058 (Table 3). Uninfected leaves had a lower starch content (mean of 1.08 μg mm^-2^) than infected leaves; however, as *C*Las exposure increased from low to high starch content decreased (means of 11.31, 5.69, 3.55 μg mm^-2^, respectively). Uninfected leaves had the lowest visual rating score, which increased as *C*Las exposure increased from a mean of 0.22 in the uninfected to a mean of 2.22 in the highest exposure group.

**Table 3.**
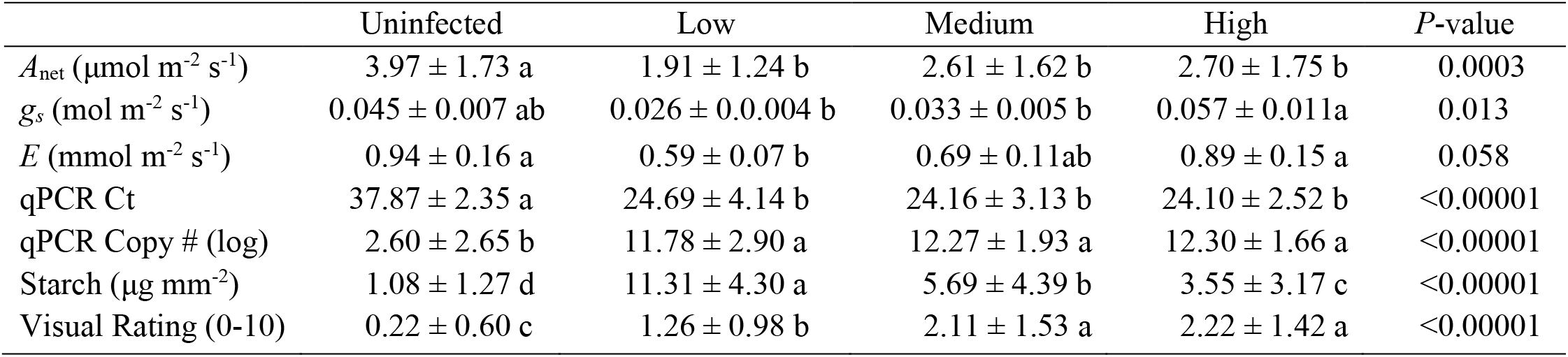
Effect of huanglongbing symptom category on leaf-level net CO_2_ assimilation (*A*_net_), stomatal conductance (*g*_*s*_), transpiration (*E*), cycle threshold (Ct) and Copy # count of *Candidatus* Liberibacter asiaticus, foliar starch content, and visual disease rating in ‘Valencia’ sweet orange (*Citrus x sinensis* [L.] Osbeck). *P-*value of analysis of variance with infection categories as the fixed effect. Means (± standard error) followed by different letters are significantly different at *P*<0.05 according to Fisher’s LSD.

*A*_net_ had a positive relationship with *C*Las exposure at the shoot-level. High exposure shoots had the highest *A*_net_ at a mean of 3.37 μmol m^-2^ s^-1^, which decreased with a shortened exposure down to a mean of 1.91 μmol m^-2^ s^-1^ in the uninfected shoots (Table 4). This pattern was similar for both *g* and *E*, with those of two or more infected exposure groups, including the high exposure group for both variables, showing higher values than the uninfected plants. However, as with leaf-level *E*, the *P-*value for exposure category differences was inconclusive at *P=*0.061 (Table 4). Reductions in mean leaf area (17.59 cm^2^ in the uninfected to 11.38 cm^2^ in the highest exposed group) and total shoot surface area (133.5 cm^2^ in the uninfected to 68.3 cm^2^ in the highest exposed group) were found as *C*Las exposure increased. Exposure did not affect internode length. Despite the different pattern of impact of *C*Las exposure, the overall range of values for shoot- and leaf-level *A, E*, and *g* were similar, except in the unexposed category, in which shoot-level values were markedly lower than at the leaf level.

**Table 4.** Effect of huanglongbing symptom category on shoot-level net CO_2_ assimilation (*A*_net_), shoot conductance (*g*), transpiration (*E*), and shoot morphological variables in ‘Valencia’ sweet orange (*Citrus x sinensis* [L.] Osbeck). *P-*value of analysis of variance with infection categories as the fixed effect. Means (± standard error) followed by different letters are significantly different at *P*<0.05 according to Fisher’s LSD.

|  | Uninfected | Low | Medium | High | P-value |
| --- | --- | --- | --- | --- | --- |
| $A_{\text{net}}$ ($\mu\text{mol m}^{-2} \text{s}^{-1}$) | 1.91 ± 1.07 b | 2.42 ± 0.88 b | 2.44 ± 0.52 ab | 3.37 ± 1.34 a | 0.032 |
| $g$ ($\text{mol m}^{-2} \text{s}^{-1}$) | 0.03 ± 0.005 b | 0.05 ± 0.007 a | 0.044 ± 0.005 ab | 0.059 ± 0.008 a | 0.039 |
| $E$ ($\text{mmol m}^{-2} \text{s}^{-1}$) | 0.45 ± 0.09 b | 0.72 ± 0.1 ab | 0.65 ± 0.08 ab | 0.85 ± 0.11 a | 0.061 |
| Stem length (mm) | 112.50 ± 22.5 | 112.30 ± 19.7 | 91.20 ± 26.1 | 90.30 ± 30.6 | 0.13 |
| Mean leaf area (cm<sup>2</sup>) | 15.79 ± 3.39 ab | 17.59 ± 5.86 a | 15.40 ± 6.33 ab | 11.38 ± 2.08 b | 0.054 |
| Shoot surface area (cm<sup>2</sup>) | 117.40 ± 42.0 a | 133.50 ± 42.8 a | 100.40 ± 48.8 ab | 68.30 ± 27.1 b | 0.014 |
| Leaves per shoot | 6.75 ± 1.49 | 7.22 ± 1.48 | 6.25 ± 2.25 | 5.33 ± 1.58 | 0.15 |
| Internode length<br>(mm) | 17.41 ± 5.49 | 15.99 ± 3.92 | 15.33 ± 3.99 | 16.99 ± 2.67 | 0.74 |

### Infection, starch, photosynthesis, and morphological variables were correlated

At the leaf level, when including uninfected trees, positive correlations were found between bacterial copy count and starch content (*r*=0.609, *P*<0.00001), *C*Las copy number and visual rating (r=0.459, *P*<0.00001), and starch content and visual rating (*r* =0.276, *P*=0.005; Appendix S6). Negative correlations were found between *A*_net_ and *C*Las copy number (*r* = − 0.322, *P*=0.0009) as well as starch content (*r*= −0.243, *P*=0.013). No correlations were found with leaf position or between *A*_net_ and visual rating. No additional correlations were found when excluding uninfected trees.

At the shoot level, when including uninfected trees, positive correlations were found between *C*Las copy number and starch content (*r*=0.716, *P*<0.00001), visual rating (*r* =0.539, *P*=0.001), and *A*_net_ (*r*=0.413, *P*=0.015); between shoot surface area and mean leaf area (*r*=0.812, *P*<0.00001) and shoot surface area and shoot length (r=0.58, *P*=0.0003); and between internode length and shoot length (r=0.344, *P*=0.046; Appendix S8). When excluding uninfected trees at the shoot level, additional positive correlations were found between starch content and mean leaf area (r=0.421, *P*=0.032) and shoot surface area (r=0.455, *P*=0.019); between shoot surface area and mean leaf area (r=0.809, *P*<0.00001) and shoot length (r=0.644, *P*=0.0004); and between internode length and *A*_net_ (r=0.558, *P*=0.003; Appendix S9).

### Linear regressions reveal additional relationships between infection and physiology

Based on the correlations we tested linear regressions of the following variables: *C*Las copy number by starch content, internode length by *A*_net_, and foliar starch content by shoot surface area. After a natural log transformation of both starch content and bacterial copy count, a positive linear relationship was found at both the leaf and shoot-level (Table 5). Further observation indicated a change in the slope as *C*Las gene copy # increased and assessment using breakpoint analysis indicated that the increased starch content at the leaf level increased at a higher rate with a breakpoint at approximately 10^11.2^ *C*Las gene copies (Figure 4A), though the shoot level data did not indicate such a change in slope (Figure 4B). A positive linear relationship (*P*=0.048) was found between foliar starch with shoot surface area when excluding uninfected shoots, though this relationship also had a low *R*^2^ (0.18) (Figure 5 and Table 5). This relationship is inverse to the *C*Las exposure duration, with the highly exposed shoots having lower starch content and lower total shoot surface area. When comparing morphological and *A*_net_ measurements, a positive linear relationship (*P*=0.003) was found between internode length and *A*_net_ when excluding uninfected shoots (Figure 6 and Table 5). These morphological adjustments are illustrated in Figure 7.

**Table 5.**
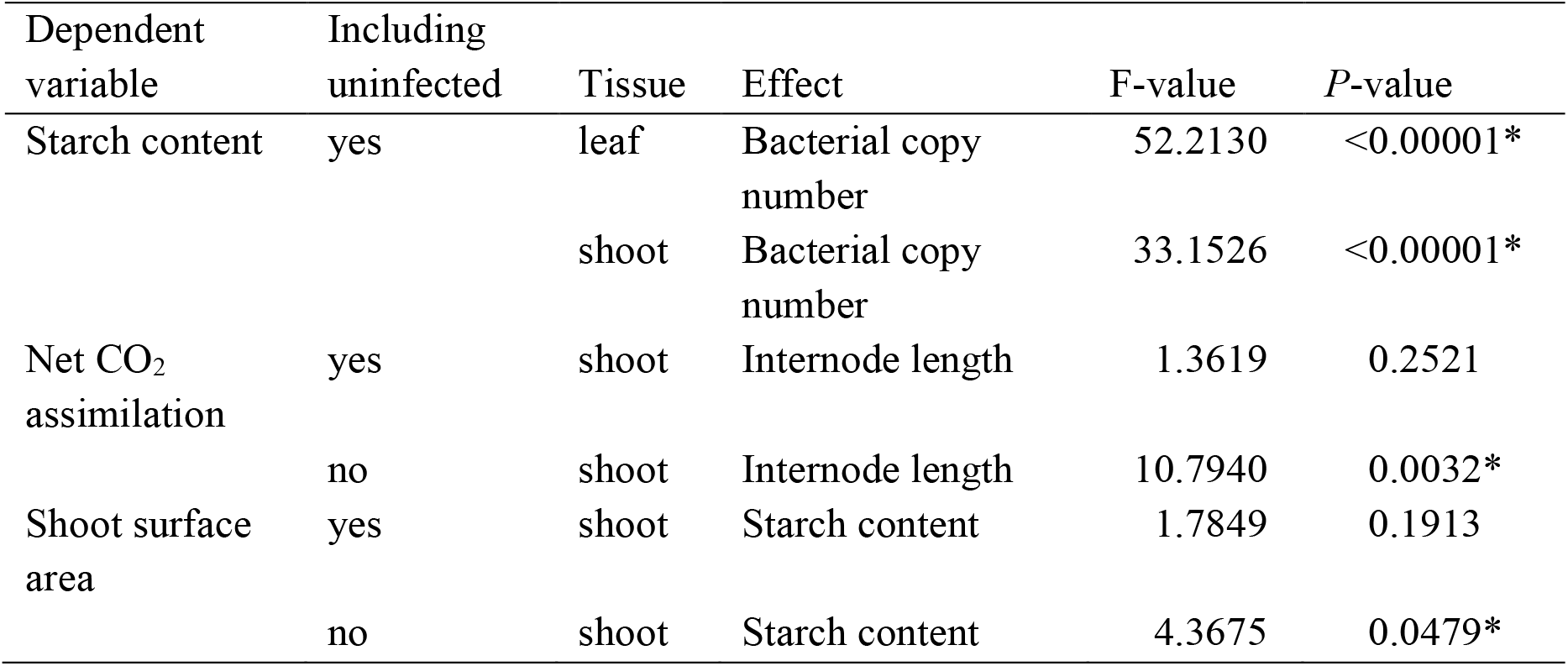
Linear regressions of effects of gene copy number of *Candidatus* Liberibacter asiaticus on foliar starch content, of internode length on net CO_2_ assimilation, and of foliar starch content on shoot surface area each assessed across three huanglongbing symptom categories including or not including a fourth category of uninfected trees, and measured at leaf or whole shoot levels. *P*-values below 0.05 are labeled with asterisks. All models included shoot nested within tree, and leaf models included leaf nested within shoot, as random variables.

**Figure 4.**
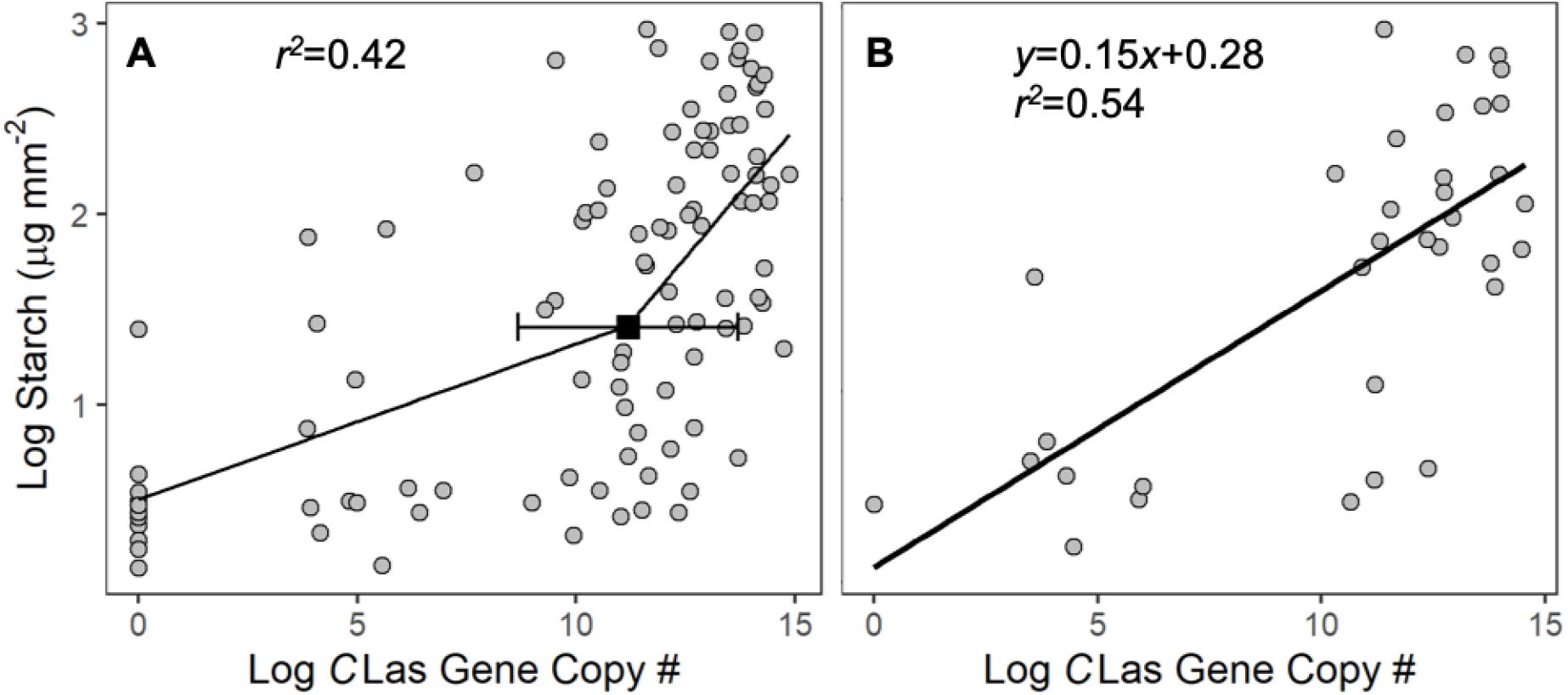
Relationship of log transformed bacterial copy count and log transformed starch content of *Candidatus* Liberibacter asiaticus-infected and uninfected ‘Valencia’ sweet orange (*Citrus x sinensis* [L.] Osbeck) A) leaves and B) shoots.

**Figure 5.**
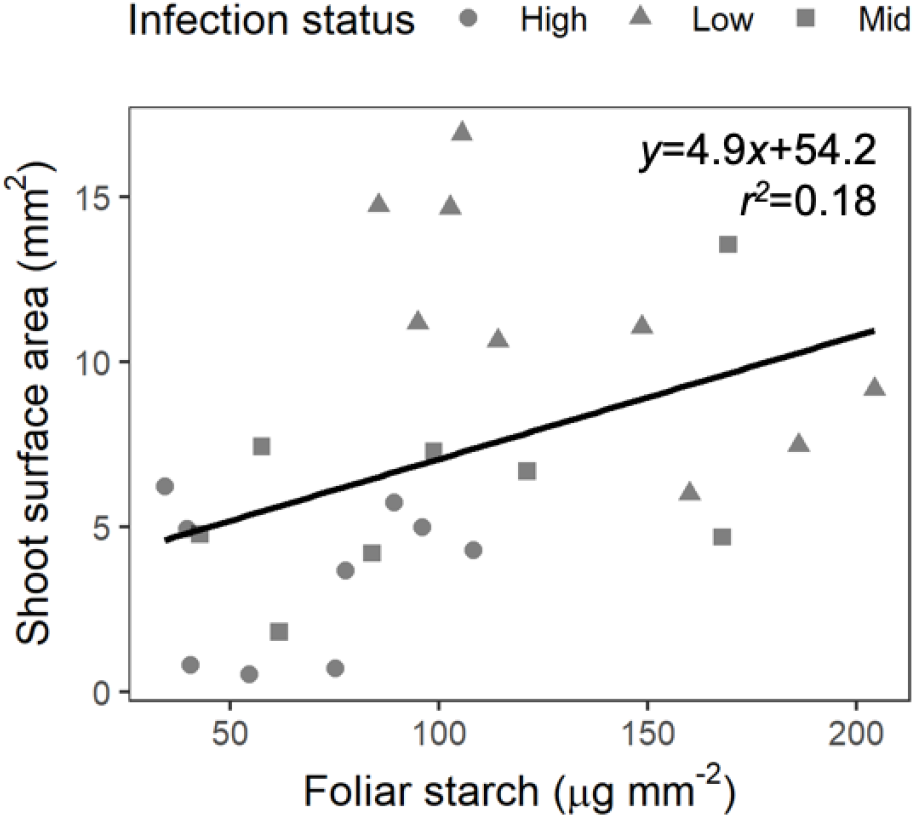
Relationship of starch and shoot surface area of *Candidatus* Liberibacter asiaticus-infected and uninfected ‘Valencia’ sweet orange (*Citrus x sinensis* [L.] Osbeck). Data point shape and color denote *C*Las-infection exposure category: Low – 1 year *C*Las-infected Asian citrus psyllid feeding and breeding, Mid – 1.5 years *C*Las-infected Asian citrus psyllid feeding and breeding, High – 2.5 years *C*Las-infected Asian citrus psyllid feeding and breeding.

**Figure 6.**
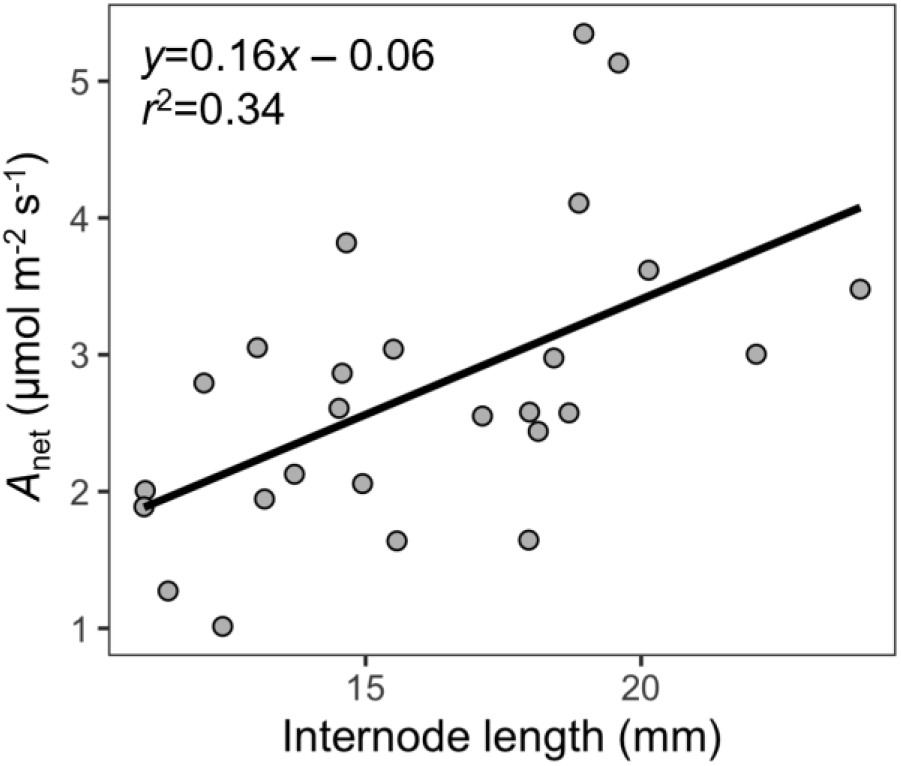
Relationship of average internode length and net CO_2_ assimilation (*A*) of *Candidatus* Liberibacter asiaticus-infected and uninfected ‘Valencia’ sweet orange (*Citrus x sinensis* [L.] Osbeck).

**Figure 7.**
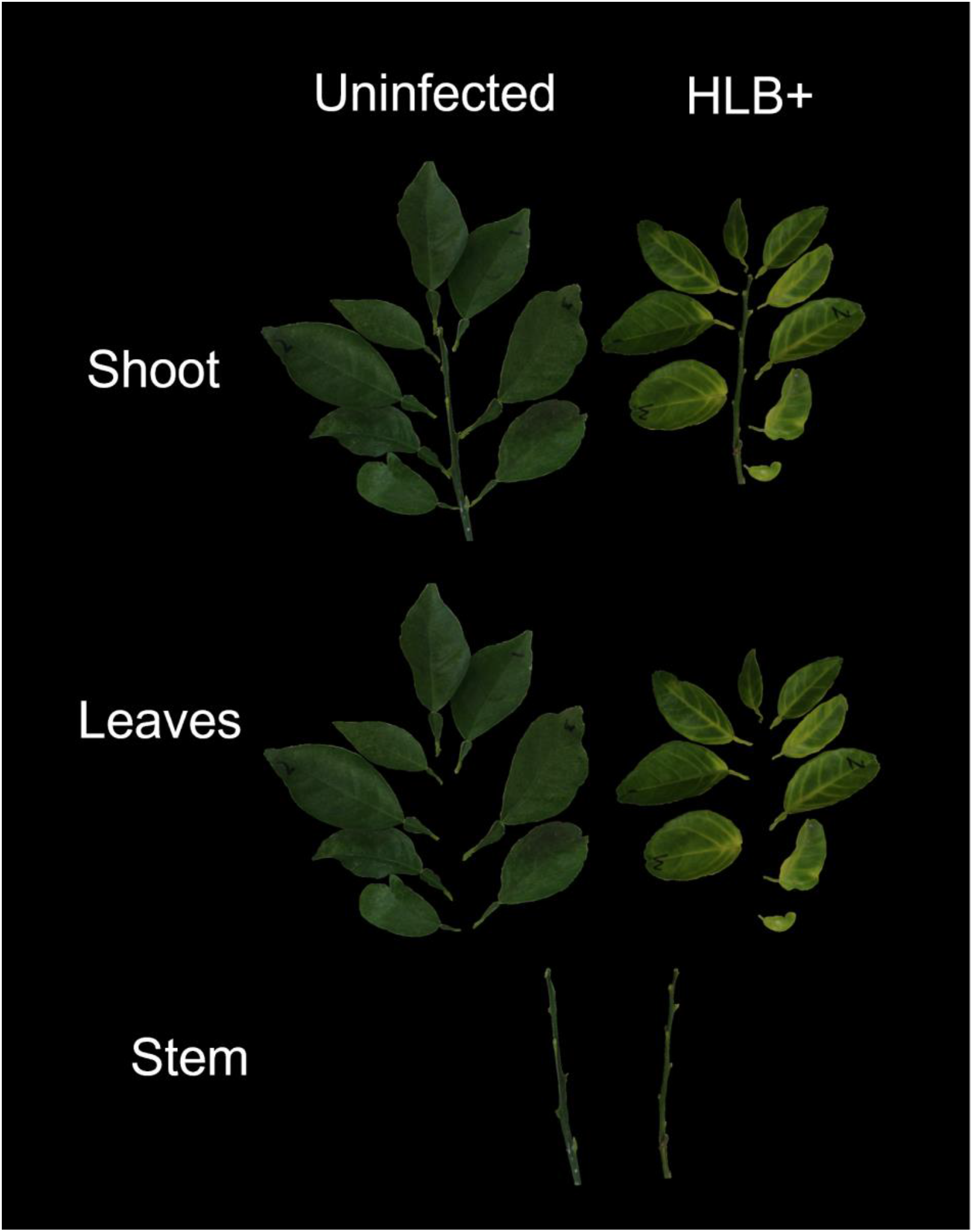
Illustration of morphological changes in citrus (*Citrus x sinensis*) induced by huanglongbing (HLB) disease caused by infection with *Candidatus* Liberibacter asiaticus.

Conductance also influenced *A*_net_ at both leaf and shoot levels, where exposure category also influenced the slope of the relationship. In leaf-level measurements medium and low exposure categories of *C*Las exposure increased the slope of the *A*_net_ response to *g*_*sw*_, while the high exposure category decreased the slope below that of the uninfected leaves (Figure 8). However, the high exposure category had more disperse data, and slope calculations may be more subject to influence of single values. At the shoot level, *C*Las infection reduced the slope of the *A*_net_ response to shoot *g*, while increasing exposure led to increasing slope.

**Figure 8.**
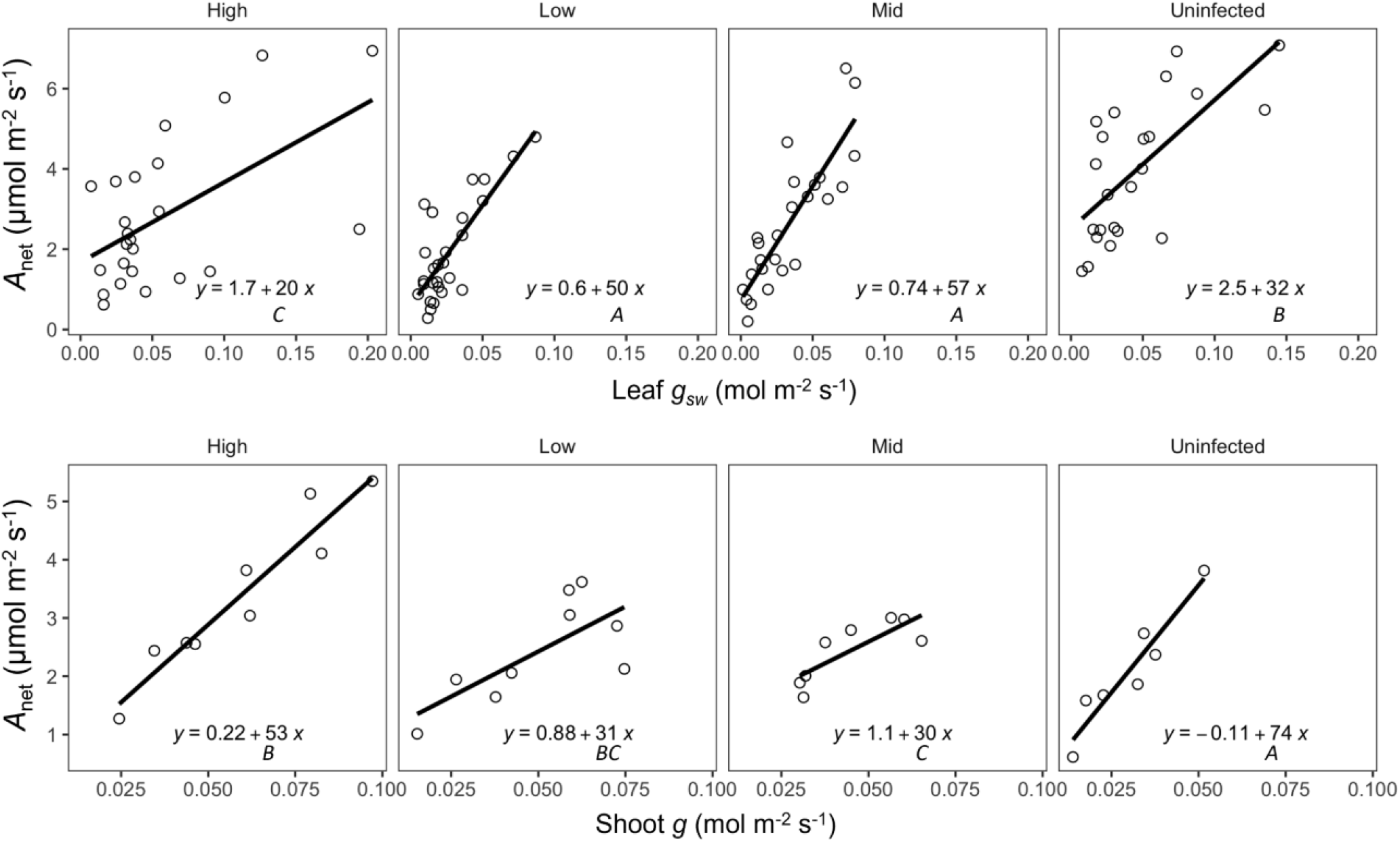
Leaf-(A) and shoot-(B) level relationships of conductance with net CO_2_ assimilation (*A*_net_) of healthy and *Candidatus* Liberibacter asiaticus-infected ‘Valencia’ sweet orange (*Citrus x sinensis* [L.] Osbeck) exposed to 1 (Low), 1.5 (Mid), and 2.5 (High) years of inoculation via Asian citrus psyllid (*Diaphorina citri* Kuwayama) feeding and reproduction. *g*_*s*_ is leaf-level stomatal conductance to water. *g* is shoot conductance to water. For both models *P* <0.0001. For leaf-level data *R*^*2*^=0.57. For shoot level data *R*^*2*^=0.80. The interaction term of *g*_*s*_ by infection category *P=*0.002; the interaction term of shoot *g* by infection category *P=*0.039. Slope values labeled with different letters are different at *P<*0.05 within leaf- or shoot-level values.

## Discussion

### Leaf and shoot light response dynamics differ

The distinct shoot light response dynamic helps to explain the success of *Citrus* in high light regions, where maximum *A*_sat_ is attained near the maximum ambient PPFD occurring in subtropical regions. The leaf-level light response curves indicating Q_sat_ occurs between 600-700 μmol m^-2^ s^-1^ PPFD agreed with earlier studies (Habermann et al., 2003). The increased light saturation point of citrus shoots compared to individual leaves could be explained by varied angle of incident radiation, as well as by partial shading of basal leaves by more distal leaves. However, *R*_d_ was much lower at the shoot level than at the leaf level. The light response curve *R*_d_ estimates are supported by the measurements at 0 µmol PAR m^-2^ s^-1^, where the estimates were similar: 1.3 ± 0.13 µmol CO_2_ m^-2^ s^-1^ for leaf and 0.47 ± 0.05 µmol CO_2_ m^-2^ s^-1^ for shoots to the estimates produced by fitting the asymptotic function. Lower *R*_d_ is an important factor in reducing the LCP. Thus, at the shoot level, citrus exhibits higher *A*_net_ at both the lower and upper ends of irradiance than is detected at the leaf level, while at PPFD ranges between 50-1000 µmol PAR m^-2^ s^-1^, shoots exhibit lower *A*_net_ than leaves. The lack of a difference in Φ is further confirmation that differences between shoots and leaves are primarily determined by factors such as morphology, not by enzymatic or photochemical factors. This might be assumed, given that some of the same leaves were involved in both measurements, though the lower rate of *R*_d_ in shoots merits further investigation. Because variations in irradiance within shoots may explain much of the difference between shoot and leaf irradiance responses, morphology is likely to have a strong impact on photosynthesis at the whole-shoot level. Thus, morphological impacts of HLB may help explain the impact of the disease on photosynthesis and growth.

### Stem assimilation contributes to whole shoot photosynthesis

Citrus vegetative stems significantly contributed to shoot *A*_app_. The stem was also more efficient in *A*_app_ on a surface area basis than leaves. The high assimilation rate of citrus stems could be explained by area-related chlorophyll content where young stems contain 50–70% more chlorophyll than adjacent leaves (Kharouk et al. 1995; Pfanz et al., 2002; Pilarski 1984; Schmidt et al. 2000; Solhaug et al. 1995). Although many factors contribute to whole-shoot assimilation, the previously unquantified contribution of the photosynthetic stem to canopy wide assimilation may be an additional factor limiting extrapolability of leaf-level gas exchange measurements to canopy-wide measurements. Stem internal re-fixation of CO_2_ in young branches of perennial woody species have been shown to compensate for 60–90% of the potential respiratory carbon loss (Pfanz et al., 2002). Although, its contributions compared to the leaves on whole shoot assimilation are relatively small, the proportion of *A*_net_ coming from the stem may depend on shoot morphological characteristics such as internode length that affect light interception in the stem. This observation highlights the importance of morphological impacts of HLB on the whole-shoot photosynthetic capacity.

### Shoot morphology and the effects of HLB of leaf and shoot photosynthesis

As with light responses, shoots and individual leaves differed in their responses to HLB. While *C*Las infection reduced *A*_net_ at the leaf level, it increased *A*_net_ at the shoot level, while inducing whole-shoot morphological adjustments over time. Although the measured values of *A*_net_ were low (∼2-4 µmol m^-2^ s^-1^), these values are typical of leaves of greenhouse-grown citrus trees (Syvertsen 1994). The morphological impacts explain the HLB effects on whole-shoot photosynthetic rates. The decreased leaf and shoot surface area effected by *C*Las exposure reflect the symptom of canopy thinning observed in infected citrus trees in the field. This process appears to occur on a scale of years. Given that most citrus leaves senesce by 17 months (Wallace et al. 1954), the changes in shoot morphology appear to occur over a complete turnover of old to new tissues.

We have not found any report in which a plant produced both longer internodes and smaller leaves in response to a disease or environmental condition. Typically, an increase in leaf size has coincided with increased internode length to reduce self-shading, such as in acclimation to low light conditions (Kavanagh, 2014; Pieters et al., 1998; Poorter and Rozendaal, 2008; Robertson, 1994), thus the peculiar morphological changes induced by HLB may be responses to the specific challenges imposed by the disease.

These specific morphological changes were associated with the shoot-level gas exchange response to HLB. Light penetration and absorption are the primary drivers of photosynthesis, and these are heavily influenced by canopy structure (Niinemets, 2007). Canopy thinning by *C*Las-induced defoliation, as described by de Graça (1991), within the high *C*Las-exposed category (fewest and smallest leaves and longest internode length) further facilitate light penetration into the canopy. With increased light penetration due to smaller leaves and greater internode length, typically shaded leaves, further from the shoot apex, and the photosynthetic stem contribute more to shoot-level assimilation, which help explain the increased assimilation efficiency found in shoots of the high *C*Las exposure group. Considering the contribution of stems to shoot *A*_app_, the relative increase in stem surface area could also contribute to the enhanced shoot *A*_net_. The increased light exposure of all parts of the shoot may also lead to compensation for the localized disease effects discussed below in reference to leaves. Although there are likely other factors that contribute to the increased assimilation efficiency of HLB-affected shoots, the correlation of these morphological variables with *A*_net_ indicates that whole-shoot morphology is a defining feature of shoot acclimation that leads to an increase in *A*_net_.

Another defining feature of the acclimation of citrus trees to *C*Las infection appears to be impacts on conductance and *E* and their interactions with *A*_net_. Citrus are low conductance species, and other studies have found strong dependence of leaf-level *A*_net_ on *g*_*s*_ under a variety of abiotic conditions, with linear responses up to *g*_*s*_ of 0.3 mol m^-2^ s^-1^, and changes in this relationship have been interpreted as relating to impacts on nonstomatal photosynthetic processes directly (Khalid et al. 2021, Shafqat et al. 2021). Depending on whether shoot or leaf levels and whether conductance or transpiration are considered, infection either initially reduced and then increased or simply increased gaseous water exchange of leaves and shoots. This may seem to contrast with the observations of reduced sap flow in HLB-affected trees (Kadyampakeni et al. 2014), but the progressively reduced shoot surface area led to a total per shoot *E* that was similar in uninfected (52.7 ± 3.78 mmol s^-1^) and high exposure infected (57.8 ± 2.78 mmol s^-1^) groups, and these instantaneous measurements do not include possible limitations to transpiration of infected plants under high diurnal evaporative demand. *C*Las infection may increase conductance by morphological (increased stomatal density) or behavioral (preventing stomatal closure) mechanisms, but the present study does not reflect on which mechanisms may be at work.

The impact of *C*Las exposure categories on the relationship between conductance and *A*_net_, however, helps clarify the impact of HLB on photosynthesis. At the leaf level, mid and low exposure categories reduced *g*_*s*_ but showed an increased relative response of *A*_net_ to *g*_*s*_. This indicates that reduced *g*_*s*_ drove the photosynthetic reduction in the lower exposure group. However, at the shoot level the impact of exposure was nearly opposite, with the low exposure group showing a marked decrease in the photosynthetic response to shoot *g*. This indicates that at the shoot level, non-conductance factors are more important, implicating carbon fixation processes directly, perhaps driven by source-sink attenuation, discussed in detail below.

The increased efficiency of *A*_net_ should not be confused with increased total photosynthesis. While the highly *C*Las-exposed shoots had higher *A*_net_ than the shoots from the low exposure group, these measures are surface-area based and do not offset carbon fixation lost in the reduction in shoot surface area with increased *C*Las exposure. Total net shoot assimilation was 40% higher for the uninfected shoots (0.032 μmol CO_2_ s^-1^) than high *C*Las-exposed shoots (0.023 μmol CO_2_ s^-1^). The shoot-level mitigation effect is evident here in that, if leaf level *A*_net_ are extrapolated to the whole shoot surface area, the estimates values would be 0.048 µmol CO_2_ s^-1^ for the uninfected shoots and 0.018 µmol CO_2_ s^-1^ for the high-exposure shoots, over- and underestimations for uninfected and infected shoots, respectively.

Although, growth is necessarily dependent on photosynthesis, numerous complexities have led to a low degree of correlation between net assimilation and growth. For example, Poorter and Remkes (1990) did not find leaf-level *A*_net_ to be correlated with relative growth rate in a comparison of numerous species, though other multi-species comparisons have found effects of net assimilation rate on relative growth rate (Li et al. 2016). In the context of growth-limiting abiotic conditions, Zait et al. (2019) found that impacts of abiotic challenges on *A*_net_ versus effects on growth rate were different among water deficit, salinity, and high temperature, identifying contributions of several components of gas exchange. Measurements of *A*_net_ using gas exchange are further complicated in that they are limited in both spatial (canopy dimensions) and time scales, and neither growth-analysis-based net assimilation rate nor gas-exchange-based *A*_net_ include the complexities of allocation and their subsequent impacts on growth (Pereira 1995). In most studies, however, total whole canopy leaf area has been found to affect growth more than photosynthetic activity on a leaf area basis (Pereira 1995, Sinclair et al. 2019). The present study provides an example of how morphological changes could overwhelm changes in *A*_net_. Despite increases in *A*_net_, *C*Las infected shoots had lower total assimilation rates because the reduction in photosynthetic surface area was greater than the relative increase in *A*_net_, leading to a reduction in assimilation per shoot. Thus, citrus plants mitigate HLB-driven losses by increasing *A*_net_ but do not increase total assimilation.

### CLas infection induces local starch and photosynthetic responses

Because *C*Las is a phloem-limited bacterium, the effects of infections on leaf carbohydrate dynamics have been the focus of much HLB-related research. Achor et al. (2010) observed starch disruption of chloroplast inner grana in association with local phloem blockage and collapse. Further, Fan et al. (2010) found increased foliar sucrose concentrations in association with starch accumulation in infected trees, where Welker et al. (*en presse*) confirmed the sucrose results and observed reduced leaf export rates. The current study suggests that these effects are localized, where *C*Las population increases induced starch accumulation at a much higher rate after reaching a population threshold at the leaf level.

Given these earlier findings regarding the impact of HLB on citrus carbohydrate relations, Cimò et al. (2013) simulated the foliar response of HLB by girdling citrus trunks, resulting in starch accumulation and a reduction in leaf level *A*_net_. This reduction had been hypothesized because of the source-sink hypothesis (Ainsworth and Bush 2011), which observes that plants downregulate *A*_net_ in response to reduced leaf carbohydrate export, a condition in which various carbohydrates may accumulate. In the case of citrus, this process was demonstrated by Syvertsen (1994), and later by Iglesias (2002) and Nebauer et al. (2011), who found *A*_net_ responded to changes in soluble sugars, which in turn also resulted in starch accumulation or depletion. Although none of the above studies addressed the condition of HLB, their interpretation of reduced photosynthesis as a response to leaf export limitation has been applied to studies of HLB, in which gene expression of photosynthetic pathways has been found to be downregulated (Balan et al. 2018). The present study found the expected reduction of *A*_net_ at the leaf level, but not at the shoot level.

The comparison of local versus systemic effects has not been addressed explicitly by studies of HLB. However, the authors of some studies have implicitly interpreted systemic effects on leaves (all studies reviewed by Balan et al. 2018) while others have hypothesized that HLB primarily affects roots (Johnson et al. 2014). The indication in the present study of a localized impact of *C*Las on leaf carbohydrate export and photosynthesis supports the interpretation of Koh et al. (2012) and Welker et al. (*en presse*), who observed callose blockage of leaf phloem as well as reduced export of ^14^C-labeled photosynthates in infected trees, while also agreeing with those of Fan et al. (2010), Rao et al. (2018), and Balan et al. (2018), all of which found alterations in carbohydrate metabolism and photosynthesis in leaves. A unique contribution of the present study is to indicate that these effects are at least partly local effects of presence of the bacterial population in the leaf, rather than strictly systemic responses to infection. One recent study found that impacts of HLB on stem transport speeds also correlated with *C*Las populations in the same stem segments, indicating an additional localized impact on transport (Welker et al. *en presse*).

Starch accumulation has been associated with *C*Las infection and has been attributed as the cause of the blotchy mottle and chlorosis symptoms seen in infected citrus leaf tissue (Etxeberria et al., 2009; Gonzalez et al., 2012; and Schaffer et al., 1986). Interestingly, we found high starch accumulation in the most recently exposed group and a relative reduction in starch content with longer exposure durations. This observation, including the recovery of the *A*_net_ response to *g* in the high exposure category indicates that the morphological acclimation leads toward an equilibrium between source area and export and transport capacity under the constraints of the disease. However, the starch levels in the high exposure group remained elevated above those of healthy trees, suggesting that a complete source-sink balance is not possible with *C*Las infection. Why photosynthesis does not further acclimate to avoid starch accumulation is an important consideration for future studies. Meanwhile, visual estimates of symptom severity increased across exposure groups, indicating that more severe symptoms were not the result of greater starch accumulation.

The *C*Las exposure categories themselves did not correspond to differences in leaf *C*Las population. Given the systemic nature of *C*Las infection, this indicates that even the mild exposure category was sufficient to attain systemic distribution of the pathogen. The lack of correlation between *C*Las titers and symptoms or disease severity has been born out in other studies, and these studies suggest wide sources of variation in titers that are not directly related to disease severity (Stover et al. 2014, 2016, Ibanez and Stelinski 2020, Vasconcelos et al. 2020). The increasing disease severity across increasing exposure groups without increases in titer indicates the cumulative chronic impact of infection, which does not appear to include increasing bacterial populations.

### Conclusion

This study provides the first direct comparison of CO_2_ assimilation between leaf and shoot levels, as well as the first comparison of assimilation in HLB-affected and healthy citrus. Shoot-and leaf-level responses were distinct. Shoot light response curves were much more gradual than those of leaves with *A*_max_ at approximately 2x the irradiance of the *A*_max_ of leaves. This was explained by the combination of leaf angles and photosynthetic stems, which contributed a mean of 10% of the total shoot *A*_app_. Likewise shoot and leaf responses to *C*Las infection differed. Where *A*_net_ was reduced at the leaf level, shoot morphological acclimation partially compensated for this downregulation in leaf-level photosynthetic activity. These morphological changes further confirm the importance of canopy density as an important measure of disease severity. This study is an example of the value of assessing physiological processes along temporal and spatial scales, with whole-shoot photosynthesis providing one tool for such efforts. Further studies should consider the impacts of HLB on photosynthetic acclimation to understand the limitations of source-sink acclimation. Understanding the nature of citrus stem gas exchange would further

## Supplemental materials

Appendix S1. Cap design of whole shoot assimilation chamber. A) bottom view – stepped drilled hole to limit low leaf interference, B) top view – straight cut to allow for stem placement, C) fit of cap into chamber, D) showing fit into chamber to prevent leakage.

Appendix S2. Straw pieces fit to cover stem of ‘Valencia’ sweet orange (*Citrus x sinensis* [L.] Osbeck) shoot to measure whole shoot gas exchange excluding stem photosynthesis.

Appendix S3. Diagram describing the measurements taken to estimate stem surface area.

Appendix S4. Standard deviation of *A*_net_ at photosynthetic photon flux density of 1000 μmols m^-2^ s^-1^ demonstrating measurement stabilization for ‘Valencia’ sweet orange (*Citrus x sinensis* [L.] Osbeck) in leaf chamber of whole shoot chamber. Dashed vertical line indicates stabilization point for leaf; dotted line indicates stabilization point for shoots.

Appendix S5. Illustration of ‘Valencia’ sweet orange (*Citrus x sinensis* [L.] Osbeck) shoots and leaves selected for to compare assimilation in leaves vs. shoots.

Appendix S6. Correlation of leaf disease and photosynthetic variables of healthy and *Candidatus* Liberibacter asiaticus-infected ‘Valencia’ sweet orange (*Citrus x sinensis* [L.] Osbeck) exposed to 0, 1, 1.5, and 2.5 years of inoculation via Asian citrus psyllid (*Diaphorina citri* Kuwayama) feeding and reproduction. Copy # - log transformed bacterial copy count, Starch – log transformed starch concentration (μg mm^-2^), Visual – visual rating of citrus greening leaf chlorosis and mottling symptoms, *A* – net CO_2_ assimilation (μmol m^-2^ s^-1^), and Position – leaf position on stem (1-n starting at apical leaf). Darkened correlation boxes denote correlations where P≤0.05

Appendix S7. Correlation of leaf disease and photosynthetic variables of *Candidatus* Liberibacter asiaticus-infected ‘Valencia’ sweet orange (*Citrus x sinensis* [L.] Osbeck) exposed to 1, 1.5, and 2.5 years of inoculation via Asian citrus psyllid (*Diaphorina citri* Kuwayama) feeding and reproduction. Copy # - log transformed bacterial copy count, Starch – log transformed starch concentration (μg mm^-2^), Visual – visual rating of citrus greening leaf chlorosis and mottling symptoms, *A* – net CO_2_ assimilation (μmol m^-2^ s^-1^), and Position – leaf position on stem (1-n starting at apical leaf). Darkened correlation boxes denote correlations where P≤0.05.assimilation (μmol CO_2_ m^-2^ s^-1^), and Position – leaf position on stem (1-n starting at apical leaf). Darkened correlation boxes denote correlations where P≤0.05.

Appendix S8. Correlation of shoot disease and photosynthetic variables of healthy and *Candidatus* Liberibacter asiaticus-infected ‘Valencia’ sweet orange (*Citrus x sinensis* [L.] Osbeck) exposed to 0, 1, 1.5, and 2.5 years of inoculation via Asian citrus psyllid (*Diaphorina citri* Kuwayama) feeding and reproduction. Copy # - log transformed bacterial copy count, Starch – log transformed starch concentration (μg mm^-2^), Visual – visual rating of citrus greening leaf chlorosis and mottling symptoms, *A* – net CO_2_ assimilation (μmol m^-2^ s^-1^), LA – mean leaf surface area (cm^2^), SA – total shoot surface area (cm^2^), Length – stem length (mm), and Inter – average internode length (mm). Darkened correlation boxes denote correlations where P≤0.05.

Appendix S9. Correlation of shoot disease and photosynthetic variables of healthy and *Candidatus* Liberibacter asiaticus-infected ‘Valencia’ sweet orange (*Citrus x sinensis* [L.] Osbeck) exposed to 1, 1.5, and 2.5 years of inoculation via Asian citrus psyllid (*Diaphorina citri* Kuwayama) feeding and reproduction. Copy # - log transformed bacterial copy count, Starch – log transformed starch concentration (μg mm^-2^), Visual – visual rating of citrus greening leaf chlorosis and mottling symptoms, *A* – net CO_2_ assimilation (μmol m^-2^ s^-1^), LA – mean leaf surface area (cm^2^), SA – total shoot surface area (cm^2^), Length – stem length (mm), and Inter – average internode length (mm). Darkened correlation boxes denote correlations where P≤0.05.

## Acknowledgements

The authors wish to thank Myrtho Pierre for assistance in gas exchange measurements, Rebecca Ebert for assistance with starch measurements, and Jason Hupp for providing the design for the adapter with which the whole-shoot chamber was developed.

## Authors’ contributions

MK, CV, and DR conceived experiments. MK developed whole-shoot chamber and conducted experiments. MK analyzed data and wrote manuscript. CV supervised experiments. CV and DR supervised analyses and manuscript writing, and revised manuscript.

## Data availability

All data available upon reasonable request to corresponding author.

