## Supplement 5 for "Citrus photosynthesis and morphology acclimate to phloem-affecting huanglongbing disease at the leaf and shoot levels"

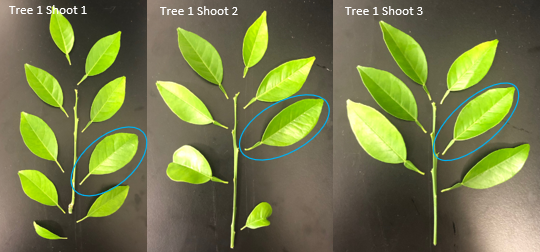

Appendix S5. Illustration of ‘Valencia’ sweet orange (*Citrus x sinensis* [L.] Osbeck) shoots and leaves selected for to compare assimilation in leaves vs. shoots.
