## Supplement 8 for "Citrus photosynthesis and morphology acclimate to phloem-affecting huanglongbing disease at the leaf and shoot levels"

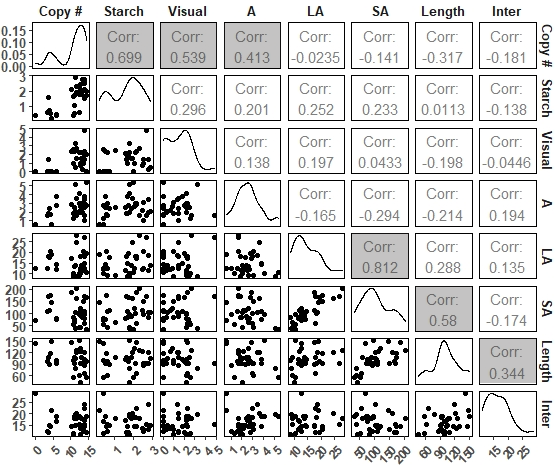

Appendix S8. Correlation of shoot disease and photosynthetic variables of healthy and *Candidatus* Liberibacter asiaticus-infected ‘Valencia’ sweet orange (*Citrus x sinensis* [L.] Osbeck) exposed to 0, 1, 1.5, and 2.5 years of inoculation via Asian citrus psyllid (*Diaphorina citri* Kuwayama) feeding and reproduction. Copy # - log transformed bacterial copy count, Starch – log transformed starch concentration (μg mm^-2^), Visual – visual rating of citrus greening leaf chlorosis and mottling symptoms, *A* – net CO_2_ assimilation (μmol m^-2^ s^-1^), LA – mean leaf surface area (cm^2^), SA – total shoot surface area (cm^2^), Length – stem length (mm), and Inter – average internode length (mm). Darkened correlation boxes denote correlations where P≤0.05.
