## Supplement 1 for "Citrus photosynthesis and morphology acclimate to phloem-affecting huanglongbing disease at the leaf and shoot levels"

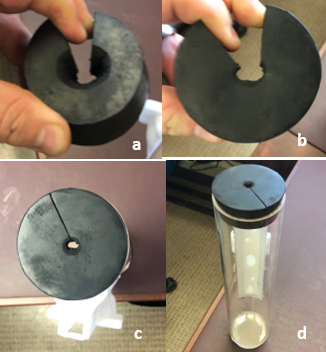

Appendix S1. Cap design of whole shoot assimilation chamber. A) bottom view – stepped drilled hole to limit low leaf interference, B) top view – straight cut to allow for stem placement, C) fit of cap into chamber, D) showing fit into chamber to prevent leakage.
