## Supplement 2 for "Citrus photosynthesis and morphology acclimate to phloem-affecting huanglongbing disease at the leaf and shoot levels"

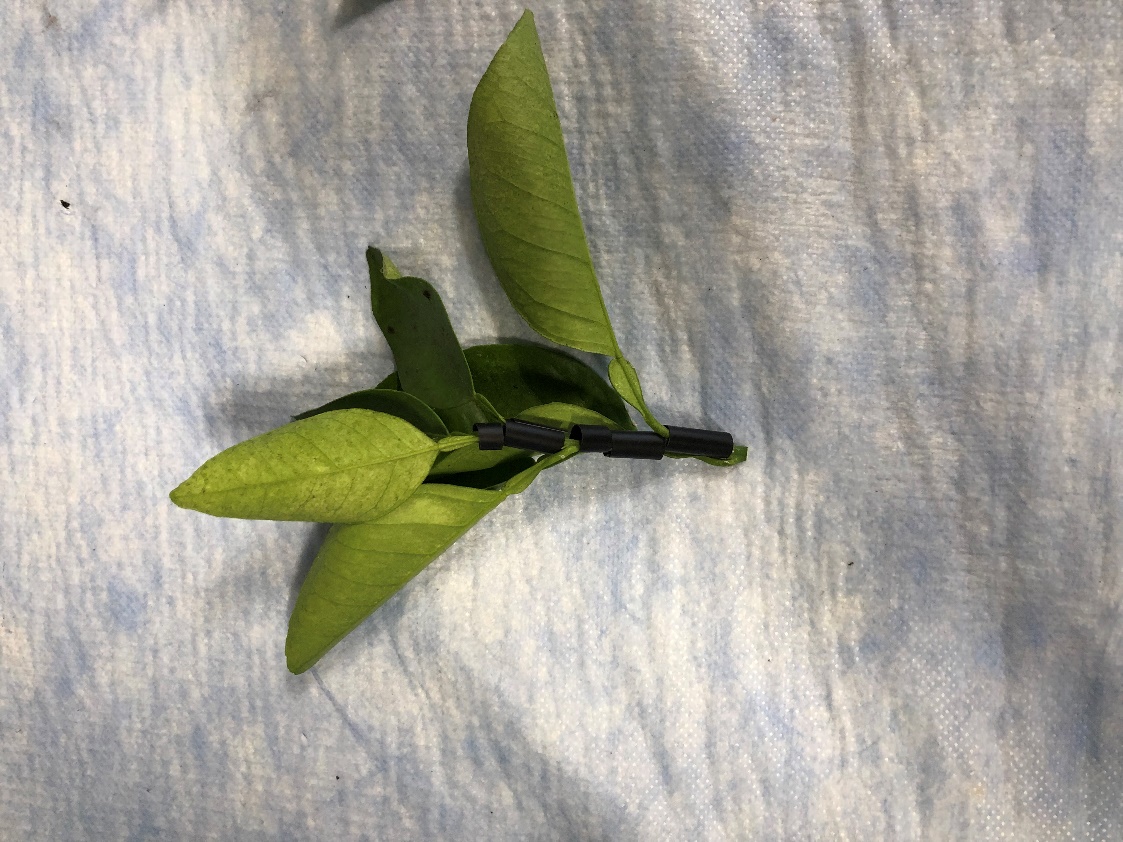

Appendix S2. Straw pieces fit to cover stem of ‘Valencia’ sweet orange (*Citrus x sinensis* [L.] Osbeck) shoot to measure whole shoot gas exchange excluding stem photosynthesis.
