## Supplementary figures and images for "Citrus photosynthesis and morphology acclimate to phloem-affecting huanglongbing disease at the leaf and shoot levels"

### Supplement 3

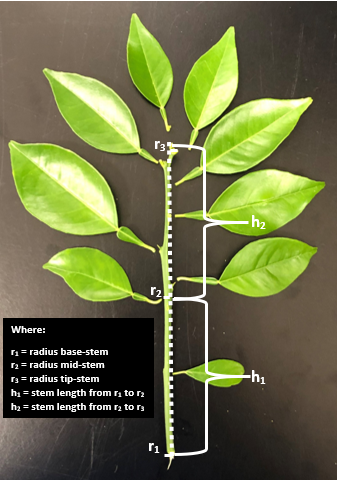

Appendix S3. Diagram describing the measurements taken to estimate stem surface area.
