## Supplement 4 for "Citrus photosynthesis and morphology acclimate to phloem-affecting huanglongbing disease at the leaf and shoot levels"

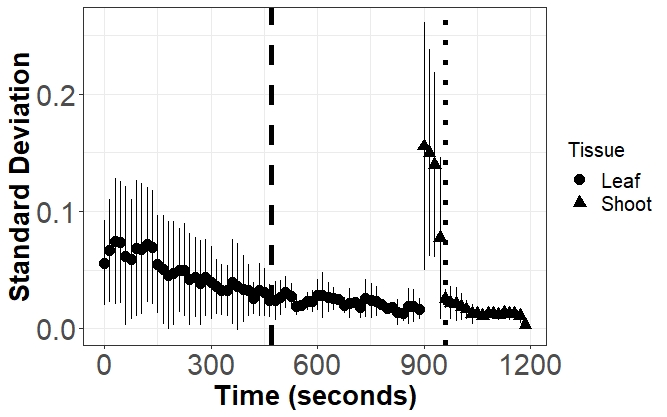

Appendix S4. Standard deviation of *A*_NET_ at 1000 μmols m^-2^ s^-1^ PPFD demonstrating measurement stabilization for ‘Valencia’ sweet orange (*Citrus x sinensis* [L.] Osbeck) in leaf chamber of whole shoot chamber. Dashed vertical line indicates stabilization point for leaf; dotted line indicates stabilization point for shoots.
